# Proximal perimeter encoding in the rat rostral thalamus

**DOI:** 10.1101/417881

**Authors:** Pawel Matulewicz, Katharina Ulrich, Md Nurul Islam, Mathias L Mathiasen, John P Aggleton, Shane M O’Mara

## Abstract

Perimeters are an important part of the environment, delimiting its geometry. Here, we investigated how perimeters (vertical walls; vertical drops) affect neuronal responses in the rostral thalamus (the anteromedial and parataenial nuclei in particular). We found neurons whose firing patterns reflected the presence of walls and drops, irrespective of arena shape. Their firing patterns were stable across multiple sleep-wake cycles and were independent of ambient lighting conditions. Thus, rostral thalamic nuclei may participate in spatial representation by encoding the perimeters of environments.

## Introduction

The position and heading of an animal within the environments it navigates are encoded by networks of neurons with differing spatial firing properties. The major components of this network are hippocampal formation place cells, which fire when the animal moves through a particular location in space (1–4); entorhinal cortex grid cells, whose multiple sharply-localized firing fields form a tessellating pattern within the environment (5, 6); and the widely-dispersed head-direction cells (HD), which fire when the animal is oriented in a certain direction (7–14). Boundaries in environments (vertical walls, vertical drops or other impassable features, such as water courses) are important features of the natural and built environment, constraining and directing behavioural trajectories. The neural representation of boundaries are therefore of interest in spatial processing (15–18). There are some previous reports of units responsive to the presence of borders or perimeters: units responsive to geometric borders have been described in entorhinal cortex (15), and to perimeters in the rostral thalamus (17). Moreover, the boundary-vector cells (BVCs) of subiculum (19–21) respond to boundaries of a specific distance and direction from the position of the animal. Here, we examine the phenotype of rostral thalamus perimeter cells: we show that neurons in the rostral thalamus (the anteromedial and parataenial nuclei, in particular) encode proximal perimeters (opaque and transparent walls, and drops), irrespective of ambient lighting conditions, and their firing is stable over multiple sleep-wake cycles. The proximal perimeter encoding of these rostral thalamic neurons may complement or support boundary coding by subicular BVCs, or geometric border coding in the entorhinal cortex.

## Results

### Unit Isolation Quality

A total of 180 well-isolated units were recorded in the rostral thalamus (see Methods: units were located particularly in the parataenial nucleus (PT), the anterior part of the paraventricular thalamic nucleus, (PVA) and anteromedial thalamic nucleus (AM)) of ten, freely-moving, rats in open-fields with fully-enclosed and partially-enclosed transparent and opaque wall configurations. Square and circular open-field environments were constructed, the former with two adjacent opaque or transparent walls at right angles, or a combination of both. With two adjacent walls present, the perimeter was defined by two vertical walls and two vertical drops. Nineteen cells (11%) showed, by visual inspection, differential activity at the perimeter of the environment. Once isolated, ‘perimeter-active’ units were recorded for several trials over at least 2 consecutive days under different environmental conditions. Particular care was taken to establish and verify each perimeter cell individually and ensure unit stability between recording trials, using the Bhattacharyya Coefficient (BC; See Methods). The BC values for similar units was 0.85 ± 0.02 (mean ± SEM). Units with similar spatial properties within the same trial, but identified on a different tetrode, were excluded from the analysed dataset to avoid inadvertent double-counting of cells, and possibly leading to a conservative under-counting of unit numbers. Despite the recording depth in the brain, units were well-isolated, and validating indices provided very low values across most recording sessions (Supplementary Fig. 1). On further checking each of the clusters, we found that they arise from conservative cluster-cutting, because corresponding enclosure ratio (ER) and L-ratio (see Methods) were very low, verifying that the clusters were consistent in capturing similar spikes.

### Firing pattern phenotypes of perimeter-tuned units

We observed three principal firing patterns among recorded perimeter-tuned units:

I. Units with increased firing when the animal explored close to the drops at the perimeter of the open field and with low-firing zones, with respect to baseline, near the walled perimeter (42%, units: 1, 5, 6, 8, 14, 16, 17, 19; Fig. 1,2)
II. Units with low-firing zones close to a drop-type perimeter and high-firing profile, with respect to baseline, close to the walls (53%, units: 2, 3, 4, 7, 10, 11, 12, 13, 15, 18; Fig. 1,2)
III. Units with a high activity and high-firing profile close to the vertical wall only (5%, unit: 9; Fig. 2).

Units were first selected for further analysis by visual inspection. Then, to determine that the units were tuned to spatial location, and did arise not by chance, we conducted a randomization analysis (described in the Methods section). Randomizing spike firing, and computing the specificity indices, revealed that spatial firing preference is not random, as the spatial coherence for recorded spike-firing is higher than the 95^th^-percentile in randomized firing. Similarly, sparsity of firing was compared to the lower limit of the confidence interval. Additionally, the location of the animal was also used to determine the contribution to unit firing. One apparently anomalous unit (#16, rotated trial; Fig. 2B(II)) did not fulfil the selection criteria for being considered as a spatially-tuned, except by visual identification. However, unit stability tests confirmed that this was the same unit as that recorded in Fig. 2B(I). Therefore, we considered this unit for further analysis.

**Figure 1.**
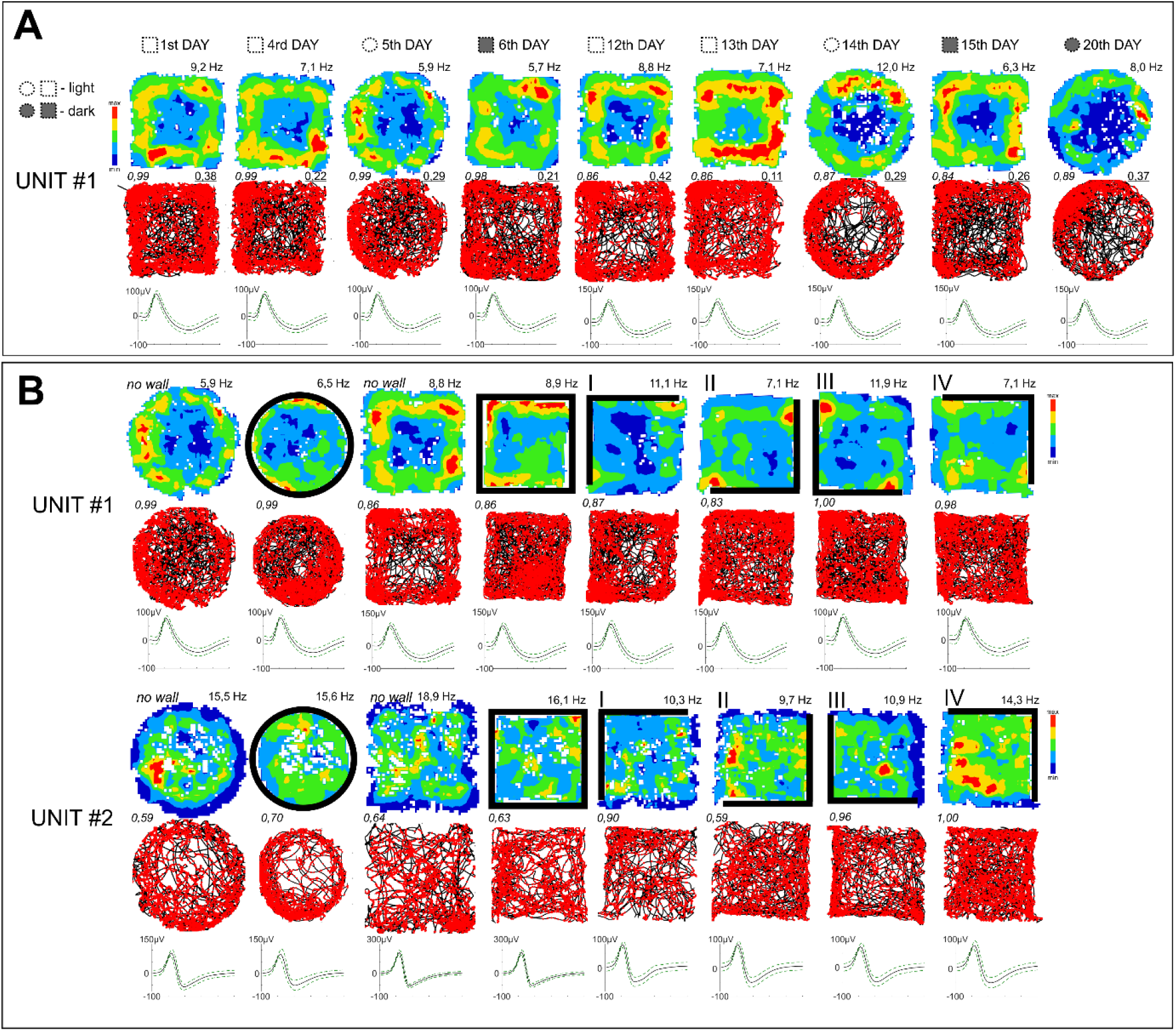
Perimeter-tuned cells recorded in the rostral thalamus over multiple days and arena shapes. A. A perimeter-tuned cell showing multi-day firing pattern stability (20 days), with firing increases at the perimeter of the environment (Category 1 units, which show increased firing when the animal explores close to the drops at the perimeter of the open field and with low-firing zones, with respect to baseline, near the walled perimeter) (elevated square; circle without walls). Light removal and changing the arena shape did not affect firing patterns of the unit or its waveform properties (unit’s similarity level). B. Representative perimeter-tuned units recorded in differing arena types. Unit 1 is the same unit as in A. Unit 2 showed is a Category 2 unit, with low-firing zones close to a drop-type perimeter and high-firing profile, with respect to baseline, close to the walls. The perimeter-tuned firing pattern followed the repositioning of environmental boundaries (solid lines represent the position of full walls). Relocation of the wall position in the partial-walled arenas in the recording arena (I-IV) affected the location-specific firing pattern of perimeter-tuned cells, but the general shape of the arena (square or circle) did not affect spatial properties of these units. Vertical walls are indicated by solid lines; transparent walls are indicated by thin lines; open arenas are unmarked. For each unit, data recorded are presented (from top to bottom): firing intensity map with a maximum firing frequency (Hz), path of the animal recorded during the recording session with superimposed firing activity and spike waveform (the solid line is the mean spike waveform and dashed lines are mean±SD of the spike waveform). Italic – unit similarity level between trials (Bhattacharyya coefficient) based on the waveform; underscore – Skaggs information content (see Materials and Methods for full description). The black lines represent the path of the animal, and the red dots the firing of a spike on that path.

**Figure 2.**
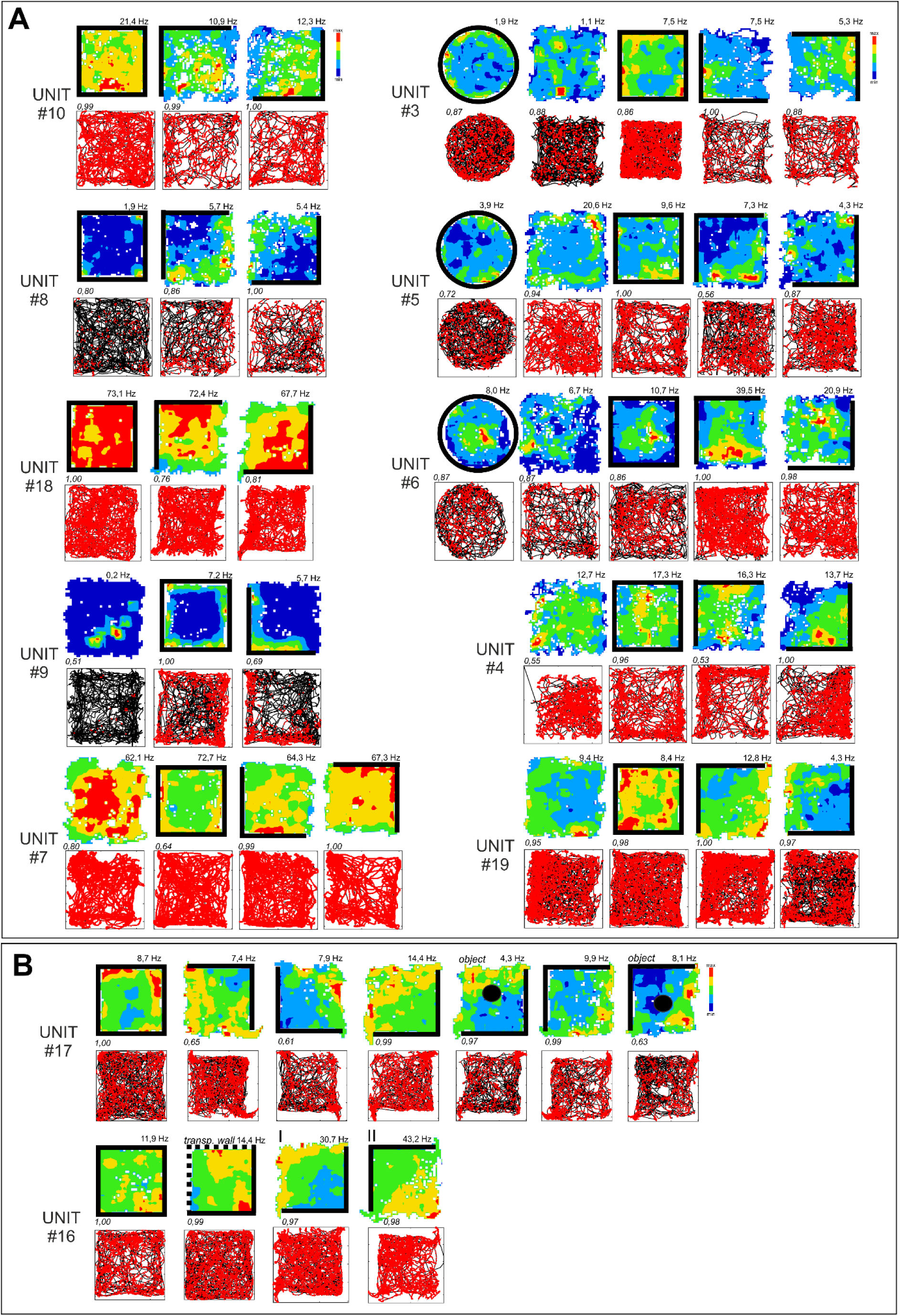
Firing properties of perimeter units recorded in different arena types. A. Firing properties of 12 perimeter units recorded in different arena types (square/circle with/without walls) and after the repositioning of the partial wall. The perimeter-tuned firing pattern followed the repositioning of the environmental boundaries (rotation of walls/no walls arena). Units with increased firing when the animal explored close to the drops at the perimeter of the open field and with low-firing zones, with respect to baseline, near the walled perimeter (42%, units: 1, 5, 6, 8, 14, 16, 17, 19) B. Firing properties of perimeter unit recorded in the presence of an object (Unit #17) or transparent walls (Unit #16). These types of environmental modifications did not affect unit firing patterns linked with the position of arena boundaries (drop-type edges in both cases). Vertical walls are indicated by solid lines; transparent walls are indicated by thin lines; open arenas are unmarked. Object is marked by the solid circle in B, fifth and seventh panels. For each unit, data recorded are presented (from top to bottom): firing intensity map with a maximum firing frequency (Hz), path of the animal recorded during the recording session with superimposed firing activity. Italic – unit similarity level between trials (Bhattacharyya coefficient). Relocations of wall position in the partial-walled arenas in the recording arena (I-II).

We compared the firing rates (of the same unit) of matched portions of the arena (either with walls or drops) over successive trials, with a 180° rotation of the partial wall for 17 pairs of trials (see Methods). We found a high correlation between rotations (0.72±0.044; Spearman’s; Fig. 4). By contrast, the correlation without rotating the arena was 0.12±0.028 (mean±SEM), whereas the correlation in non-spatial related units was 0.17±0.1 (mean±SEM). The firing patterns of these units are thus tuned to the orientation of the partial wall.

### Changing arena shape does not affect perimeter-tuned cells

The position of the perimeter affected the location-specific firing pattern of perimeter-tuned cells, but the general shape of the arena (square or circle) did not affect spatial firing pattern by these units. A clear effect on the unit’s firing frequency was observed depending on the relative positions of the walls or drop-edge in the environment (a 20-80% change in the frequency range). The perimeter-tuned firing pattern followed the relocation of the wall position in the partially-walled arenas (Fig 1, 2, 3).

### Perimeter-tuned cells are stable across days

The firing patterns of spatially-tuned cells are stable across days. Sixteen cells were recorded across at least two consecutive days, and three were stably recorded for between five and twenty days (Fig. 1, 2). To verify that the same units had been recorded between trials/days, we used cluster similarity measurement techniques (see Methods) to determine whether the units are stable across days.

### Part-removal of walls causes some perimeter cells to remap

In some experiments, we initially recorded perimeter cells with all four walls present (Fig 2). Subsequently, we removed two adjacent walls, leaving two walls intact at a right angle. Initially, these cells fired adjacent the whole four-wall perimeter (when all four walls were present; see Fig 2A, Unit #10, panel 1 for an example). Removing two walls resulted in an increase of firing adjacent the vertical drops, and a decrease in firing adjacent the remaining walls (when just two adjacent walls were present; see Fig 2A, Unit #10, panels 2 and 3, for an example).

### Lighting conditions do not affect perimeter cells

Changing visual conditions from light to dark did not affect the perimeter-tuned units. The firing patterns of units recorded in the same arena type in the light and dark were similar (Fig. 1), indicating that perimeter units are not affected by ambient lighting conditions.

### Perimeter-related firing is not directionally-tuned

Perimeter-tuned units are not directionally-tuned, and perimeter units do not show any significant relationship between perimeter responsivity and directional tuning (Rayleigh’s z-test). Among all of the other recorded cells (non-spatial related units), we observed 7 head directional units. The width of the receptive field and resultant vector length for these units were 64.57±5.54 and 0.39±0.042 (mean ± SEM) respectively.

### Angular head velocity and speed do not affect firing of perimeter units

There was no correlation between the speed of animal movement and perimeter unit firing rate (Skaggs Information Content for running speed vs spike rate is 0.080±0.009; Pearson’s R and P values are 0.19±0.06 and 0.16±0.03). Similarly, the Skaggs information content for angular velocity vs spike rate is 0.11±0.012 and the Pearson’s R and P values were 0.15±0.04 and 0.34±0.04 respectively.

### Temporal and waveform characteristics of perimeter units

The firing rate of the units was within 6.3-12.6Hz (95% CI). The mean spike amplitude, the average spike width, and the mean height of the units were 110.3±3.9 μV, 201.5±5.1μs and 158.5±4.67 μV respectively (mean ± SEM) (see Table 1 for details). We did not find the characteristic alternating peaks and troughs in the autocorrelation histogram of the inter-spike intervals of theta-modulated units across recorded perimeter cells, with one exception (unit# 6), where clear modulation of firing activity in the theta frequency range was observed.

**Table 1.** Electrophysiological classification of thalamic perimeter units (mean±SEM).

|  |  |
| --- | --- |
| N | 19 |
| Mean spike amplitude | 110.3±3.9 μV |
| Average spike width | 201.5±5.1 μs |
| Mean height | 158.5±4.67 μV |
| Mean frequency | 9.45±1.58 Hz |
| Maximal place map frequency | 13.71±1.53 |
| Spatial coherence | 0.44±0.015 |
| Spatial information content (Skaggs) | 0.52±0.052 |

### Intra-arena objects and transparent walls did not affect unit activity

There was no effect on the firing pattern of the perimeter-tuned cells after the introduction of transparent walls into the recording arena. Discrete objects introduced into the environment did not change the firing properties of perimeter units, which maintained a perimeter-tuned firing pattern despite the presence of the object in the recording arena (Fig. 2B).

### Histological analysis

Histological verification of recording sites, based on the visible electrode track, supported by the known lengths and excursions of the electrodes and a standardised estimation of rostral thalamic nucleus positions from histological atlases show that recording tetrodes were positioned in rostral thalamic nuclei – the parataenial nucleus (PT), the anterior part of the paraventricular thalamic nucleus (PVA, units: #17 and #18– See the Supplementary File), as well as in the anteromedial thalamic nucleus (AM), close to the border with the anteromedial thalamic nucleus (Fig. 5).

## Discussion

We describe here a population of spatially-tuned cells in the rostral thalamus, whose firing pattern is strongly and selectively linked with the position of the geometric perimeters of the environment (opaque or transparent vertical walls or drops). The majority of units showed non-zero baseline firing within the whole arena, in marked contrast to hippocampal units, which tend to be silent outside of their receptive fields. Three major classes of recorded perimeter-tuned units could be distinguished: units exhibiting high-firing activity when the rat explored close to the environmental perimeter (walls or drops); units with low-firing activity near the perimeter (walls or drops), and units with high location-specific activity and a high-firing profile close to the walls or drops. Units preserved their perimeter-specific firing properties across days, in light and dark conditions, as well as in arenas of different shape (circle or square).

These rostral thalamic perimeter-responsive cells are not driven purely by visual inputs, because these units are stable in light and dark conditions (Fig. 1A), as well as in the presence of transparent walls (Fig. 2B). This suggests that tactile cues might play a crucial role – which in a purely tactile, wall-responsive unit, might be correct (e.g., Unit #9). However, we found units sensitive to perimeters defined by drops – and therefore lacking the somatosensory inputs deriving from the presence of the walls. Furthermore, discrete objects introduced into the environment with a different texture to the walls did not change the firing properties of perimeter units. Perimeter-tuned firing patterns are maintained, despite the presence of the object in the recording arena. Thus, there appears to be no predominating sensory information defining the presence of the perimeter of the environment, and, presumably, the receptive fields of these units are multisensory in nature. Interestingly, during the same recording trials, we observed units with an anti-perimeter-tuned firing pattern close to each other (Fig. 3). Two such units were recorded on the same tetrode, suggesting very close physical distance between each other. This proximity suggests the existence of clusters, analogous to the suggestion of clusters of HD cells (29), composed of perimeter-tuned units with opposite firing pattern properties.

**Figure 3.**
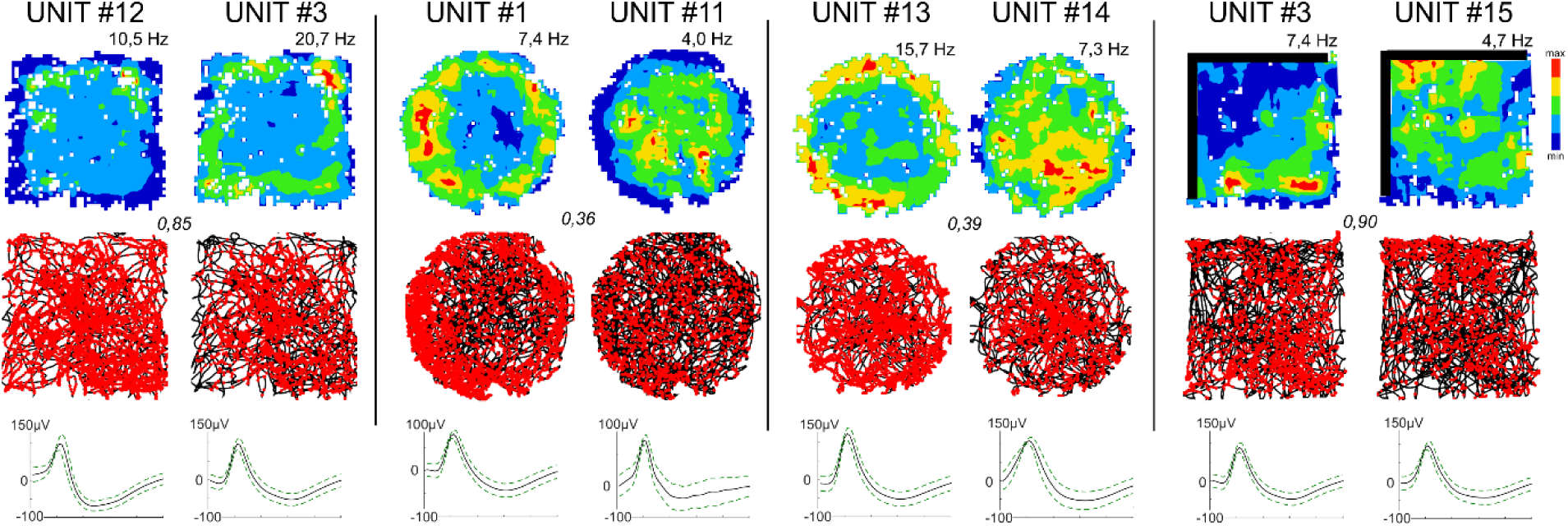
Comparison of perimeter unit pairs with remapped anti-perimeter firing patterns recorded during the same session. Four pairs of simultaneously-recorded perimeter units with an “inverse” firing pattern at the perimeters of the arena. Units #12/#3/#15 were recorded on different tetrodes, and are characterized by a high level of similarity (Bhattacharyya coefficient ≥ 0.85). Units: #1/#11 and #13/#14 were recorded on the same tetrode and are characterized by a low level of similarity (Bhattacharyya coefficient ≤ 0.39). Vertical walls are indicated by solid lines; open arenas are unmarked. For each unit, data recorded are presented (from top to bottom): firing intensity map with a maximum firing frequency (Hz), path of the animal recorded during the recording session with superimposed firing activity. Italic – unit similarity level (Bhattacharyya coefficient).

We use the phrase ‘perimeter’ units to distinguish them from the boundary-vector cells (BVCs), previously described in the subiculum (20, 21), and the border cells of entorhinal cortex (15). BVCs respond preferentially to an environmental boundary of a particular distance and direction from the rat. By contrast, the perimeter cells we describe here are characterised by their adjacency to the perimeter, and their independence from direct somatosensory input. They may, therefore, more closely resemble the border cells of the claustrum (18). These similarities include: lack of responsiveness to dark/light conditions; predictable, stable field repetition in response to both types of perimeter (walled or drop-type); no directional tuning; lack of specific sensory information defining the presence of a perimeter and occurrence of perimeter-related firing inhibition (similar to perimeter-off cells in subiculum; 21). Both nuclei share substantial reciprocal connections with entorhinal cortex and subiculum (22 – 28); perimeter units may therefore support the computation of subicular BVCs and/or entorhinal cortical border cells.

Using our conservative unit isolation criteria (see Methods), we suggest that perimeter coding is sparse (c. 10% of recorded rostral thalamic units), but similar in proportion to border cells found in entorhinal cortex (c. 10%), but less than the proportion recorded in a ‘restricted’ portion of subiculum (c. 19%-24% of units). Restricted or minor changes in electrode placement, however, seemingly result in the complete absence of subicular BVCs: compare Fig 4 of Lever et al (20) to Fig 11 of Brotons-Mas et al (36), so upper-bound estimates for proportions of subicular BVCs are uncertain. The rostral thalamic perimeter cells described here may also reflect direct subicular/cortical inputs; one possible source are the ‘annulus’ cells described by Weible et al (37) in anterior cingulate cortex (comprising 11/281 (c.4%) of recorded units), which fire adjacent the vertical boundary of an enclosed, circular arena. Alternatively, these thalamic perimeter cells may provide elemental spatial inputs to assist hippocampal and parahippocampal spatial computations, and perhaps cortical spatial processing. Finally, they might represent a component of a parallel system (together with thalamic HD cells, place cells and object cells) that is not hippocampally-or cortically-driven, which provides an autonomous coding of the position of an animal in the environment. These, and other data (17, 18) suggest that contemporary models of spatial processing may need extension to account for spatial computations that are apparent in other brain regions (38).

**Figure 4.**
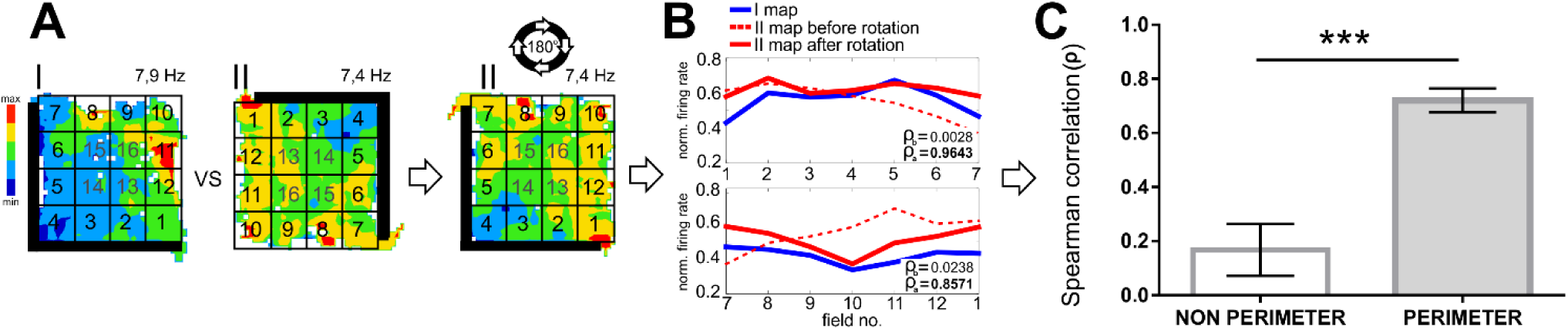
Analytical steps to evaluate the remapping of perimeter units firing with respect to environmental boundaries. A. The entire arena was re-pixelated into 4×4 squares, and each pixel was labelled from 1 to 16 as if the numbers follow a spiral starting from one end of the wall (I). For a trial where the wall was rotated by θ degree compared to the base trial, the entire map was rotated by θ to align the labels along with the base trial (II); B. The normalized mean firing rate of the new pixel was compared one to one before and after rotation. In some of the units, the firing pattern was aligned along the walls, and in others it followed the border of the arena without walls. To increase the power of the comparison between trials of the same units, the portion of the arena where the unit had been firing was split along the boundary from the other half, the corresponding normalized mean firing rates were compared using Spearman’s rank correlation coefficient (ρb and ρa for before and after rotation); C. The distribution of the coefficients in non-perimeter and perimeter units is shown in the bar plot. The data are presented as mean ± SEM. Asterisks indicate the differences between groups (Student’s t-test for independent samples; ***p ≤ 0.001).

**Figure 5.**
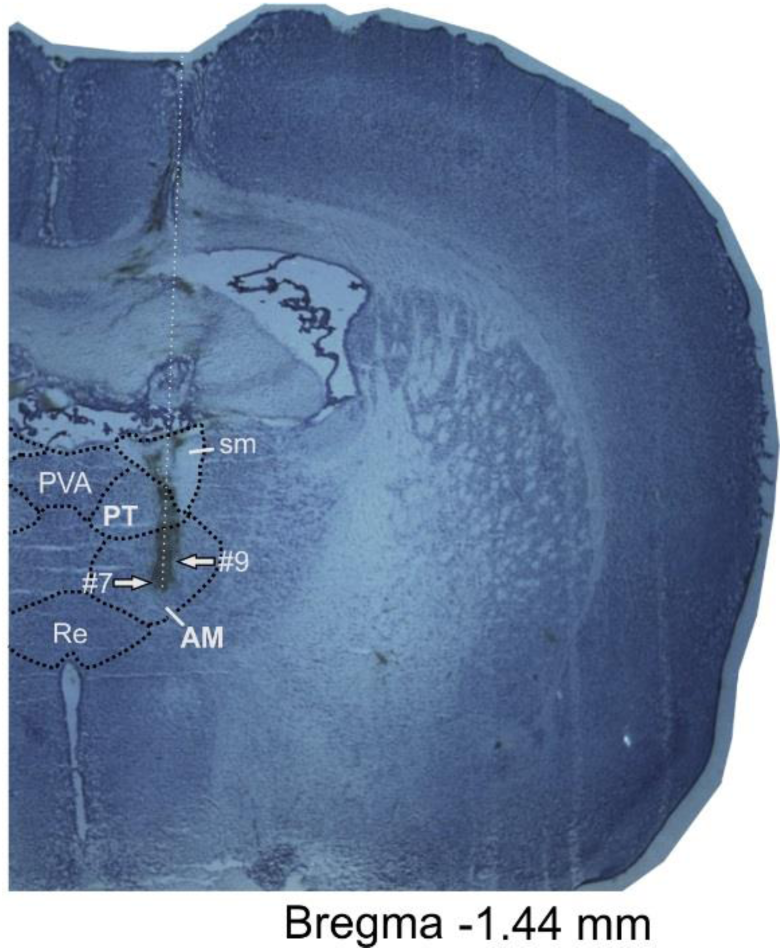
Representative histological specimen showing electrode track. Histological slide with path of the recording tetrode. Arrows depict estimated location of recording electrodes tip for unit #7 and #9. Abbreviations: AM, anteromedial thalamic nucleus; PT, parataenial nucleus; PVA, paraventricular thalamic nucleus, anterior part; RE, nucleus reuniens; sm, stria medullaris of the thalamus.

## Materials and Methods

### Animals

Experiments were performed on 10 (5–8 months) male Lister-Hooded rats (Harlan, UK) weighing 360–500 g. Upon arrival, animals were housed individually and handled by the experimenter daily for a week before being trained in the pellet-chasing task and before surgical procedures. During the stereotaxic surgery, rats were implanted unilaterally with a bundle of eight tetrodes of ø 25 μm platinum–iridium wires (California FineWire, CA, USA), mounted onto drivable 32-channel microdrives (Axona Ltd., UK) and targeted at the rostral thalamic nuclei (PT, AM, PVA). The coordinates were as follows: 1.20 mm posterior to bregma, 5.30 mm below the brain surface and 1.00-1.20 mm-1.20 mm lateral to the midline at angle of about 4.0 degrees. After the surgery, rats were allowed at least one week of recovery. Animals were then accustomed to the recording room (partially curtained room with the recording arena in the centre) and experimental procedures (1 week). During recording sessions, tetrodes were lowered very slowly through the brain (maximal rates 50 μm/day, with a total distance of 1.2 mm of tissue penetration) typically over a period of weeks to ensure a low risk of tissue damage, given the relative inaccessibility of these deep brain regions. During recordings, animals were food-restricted to 85% of their *ad libitum* body weight, and kept in a temperature-controlled laminar airflow unit and maintained on a 12-h light/dark cycle (lights on from 08:00 to 20:00 hours).

### Ethical and Regulatory Review

All experiments and experimental protocols underwent full ethical and procedural review by the Trinity College Dublin Animal Welfare Committee prior to conduct. Experiments and experimental protocols were reviewed, authorised, licenced and carried out in strict accordance with the ethical, welfare, legal and other requirements of the Health Products Regulatory Authority regulations, and were compliant with the Health Products Regulatory Authority (Irish Medicines Board Acts, 1995 and 2006) and European Union Directives on Animal Experimentation (86/609/EEC and Part 8 of the EU Regulations 2012, SI 543).

### Experimental design

In all animals, single unit activity as well as local field potential (LFP) were recorded daily (between 09.00 and 18.00) in the open field environment. During each recording session (1h), rats performed a pellet-chasing task: 20 mg food pellets (5TUL formula, TestDiet, St Louis, USA) were thrown in the arena at random locations. During each 1 hr session, animals were placed in the different arena types (15 – 30 min/arena), in light and dark conditions (randomized order). The arenas used were placed on an elevated platform (80 cm high; therefore, an 80 cm drop) and comprised of: circle with full walls (diameter 96 cm, 40 cm high)/circle without walls; square with full walls (length of 60 cm, 40 cm high)/square with transparent walls (two of four)/square without walls; square with partial walls (with differing positions of the walls) and partial walls with object present.

## Data analysis

### Unit Isolation

For off-line signal analysis, proprietary Axona software, a custom-written suite of Matlab scripts (NeuroChaT©) and GraphPad Prism 6 (GraphPad Software, La Jolla, CA, USA) were used. We used manual cluster cutting technique using TINT to isolate the activities of single-units using their waveform features (amplitude, peaks, troughs, etc.), with a criterion of a clean refractory period (≥ 2 ms) in the inter-spike interval (ISI) histogram. Similar waveforms were first grouped together using k-means clustering. The resulting clusters were tested for over-clustering, and merged if necessary. Clusters were further cleaned by manual cluster cutting.

To evaluate over-or under-clustering, and mixed multiple unit/single unit activity, we used peaks, troughs, and the first two principal components of each waveform in the channel where they were recorded most clearly (sum of square of waveform samples). We formed clusters from the tags of the unit waveforms assigned during clustering. The evaluation was accomplished using the following techniques. We calculated the Bhattacharyya coefficient (BC), which gives an approximate measure of degree of overlap between two distributions, as quantified using Equation-1(a) and 1(b). We also calculated the enclosure ratio (ER), which is the percentage of number of data points in a cluster with a higher probability of belonging to another cluster. Then, to compare cluster c1 and c2, where we assume cluster c1 is the smaller, the distance of all the data points of c1 were measured from the c1 cluster and the c2 cluster. As these distance values form a chi-square distribution, we evaluated the corresponding probability for each such distance measure using appropriate degrees of freedom. The percentage of cluster points in c1 that have a higher probability for c2 than c1 was defined as the enclosure ratio. A higher BC and lower ER will indicate that that the cluster was over-cut to make it cleaner rather than combining spikes from multiple units into one cluster. Finally, we calculated the L-ratio, derived from the sum of the probabilities that each spike that is not a cluster member is actually part of the cluster (31), divided by the total spikes in the cluster. In the original paper (21), all the spikes not part of the clusters were used for the calculation, we calculated the L-ratio in pairs of clusters, and chose the highest value as representative L-ratio measure for isolation quality.

### Spatial Properties

Once well-defined neuronal signals were isolated, they were further analysed if the rats explored the arena sufficiently (i.e. rats had to explore at least 90% of the arena in either session to be included in analysis to allow reliable calculation of spatial characteristics). The firing properties of the unit in relation to spatial (i.e., location, head direction etc.) and non-spatial stimuli (i.e., speed, angular head velocity) were calculated using temporal averaging of counted spikes. The spatial firing map was constructed by dividing the arena into pixels of 3cm×3cm, and normalizing the number of spikes that occurred in specific pixel coordinates by the total time the animal spent in that coordinate. The resultant map was smoothed by a 3×3 moving average filter. Similarly, the directional firing for each cell was obtained by dividing the entire horizontal plane into 5° bins, and dividing the total number of spikes in a bin by the total time the animal spent in that bin. Rayleigh’s Z-test was conducted to test for the non-uniformity of the head-directional tuning rate with the null hypothesis that there is a sample mean direction. Units with p<0.05 were considered as tuned to head-direction. The width of the receptive field was taken as the half-width of the tuning curve. The resultant vector length for the head-direction vs firing rate was also calculated.

Spatial coherence is the spatial auto-correlation between an unsmoothed firing map, and the map after it is smoothed. The firing map was smoothed using a box filter of length 3×3, which implies the firing rate in each pixel in the smooth map is the average of the firing rate of eight nearest neighbours and itself (32). Therefore, spatial coherence measures the extent to which firing rate in each pixel is predicted by its neighbours, and it estimates the local orderliness of the spatial firing pattern. Spatial sparsity measures the compactness of the firing field relative to the recording apparatus. It was calculated per the methods described in (33). Skaggs information content, a measure of specificity of spike-firing in the recording arena, was also calculated using the method described in (30).

To ensure that the observed firing of a spatially modulated unit is not happening by chance, we have randomized the time when the unit have fired and recalculated the firing rate map and corresponding spatial selectivity measures i.e. Skaggs information content, spatial coherence, and spatial sparsity. We repeated the procedure for 1000 times for each unit, and the 95% (5% for sparsity) percentile of their distribution was compared to the corresponding value in the original unit firing time.

We conducted multiple regression analyses to evaluate the contribution of environmental variables on the firing properties of the unit. Multiple regression represented the instantaneous spike-rate as a linear combination of environmental variables, using location, running speed, angular head velocity, head direction of the animal and its distance from the border in the arena as the environmental or independent variables. As the spike rate of the unit is non-linear in nature for head-direction, place and border cells, we used the firing rate of the unit at a particular values of the independent variable to be representative of the corresponding variable value in the regression analysis (34). We then calculated the partial correlation or the percentage of variance of the momentary firing rate explained by each variable. We repeated the procedure 1000 times on random samples with equivalent 2min long data and each sample having a span of 100msec (subsampled). The mean of the partial correlations, represented as percentage of the overall correlation, was then observed.

### Assessing the effect of running speed and angular velocity

To examine how running speed affects unit firing properties, raw speed data were smoothed using a moving average filter with a rectangular window of duration 100ms (five samples). The number of times the rat moved at a certain speed was measured using 1ms bins up to 40ms/sec (the maximum speed analysed). Spike rate was calculated by dividing the total number of spikes belonging to each running speed bin by total time the rat stayed at that speed. Speed bins at which rats did not travel for at least 1sec were then excluded. The remaining firing rates were then fitted to a linear regression equation with respect to speed bins. Skaggs information content was also calculated (30).

To calculate angular head velocity, head direction was first smoothed using a moving circular mean filter over a 100ms rectangular window. The angular velocity at time t(i) was then calculated as the slope of the linear regression line passing through head direction at t(i-2) to t(i+2) time samples after adjustment of circular zero-crossing (10). The –ve and +ve angular head velocities were regard as clockwise (CW) & counter-clockwise (CCW) angular velocities respectively. For each of these groups, the spike rate was calculated using a similar approach to that of running speed. Bin size and maximum angular velocity were set as 10deg/sec & 500deg/sec respectively. Bins with trailing zeros at higher angular velocities and those not visited for at least 1sec were removed. Both the CW & CCW spike rates were separately fitted to a linear equation, and quality of fit was assessed using a Pearson’s correlation coefficient between the fitted spike rate and the raw rate.

### Examining the similarity of units

To examine unit stability, and to track units day-to-day or trial-by-trial, their similarity was examined using cluster-quality measuring techniques. Amplitude, waveform shape and the first two principal components of the waveforms were considered as the characteristics of the units. The similarity between two comparing units were then calculated using the Bhattacharyya Coefficient which measures the amount of overlap between two normal distributions. The closed form expression for the analysis is given in Equation-1(a) and 1(b).

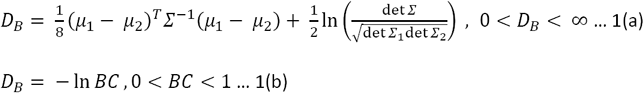

Here, D_B_ is the Bhattacharyya distance, μ _i_ and Σ _i_ are the means and the covariances of the distribution, and Σ is the average covariance of the two distributions. The Bhattacharyya Coefficient (BC) is then given by Equation 1(b). Units were compared only between the ones from same tetrode across trials (to test for similarity) or within trials (to test for dissimilarity). Units were considered similar if BC≥0.5 as the value of BC between clusters of the same electrodes within trials are 0.43 ± 0.04 (mean ± SEM).

### Comparing rotational remapping of firing maps

To observe how spatial firing remapped due to rotation of the half-walls in the arena, we devised the following analysis (Figure 4a). The entire arena was re-pixelated into 4×4 squares, and each pixel was labelled from 1 to 16. For a trial where the wall was rotated by θ degrees compared to the base trial, the entire map was rotated by θ to align the labels along with the base trial (Figure-4(a), II). After the alignment, the normalized mean firing rate of the new pixels was compared one-to-one. In some of the units, the firing patterns were aligned along the walls, while in others it followed the perimeter of the arena without walls. To increase the power of the comparison between trials of the same units, the portion of the arena where the unit had been firing was split along the perimeter from the other half. The corresponding normalized mean firing rates were compared using Spearman’s rank correlation (ρ) coefficient (Figure 4c).

### Perfusion and histology

After completion of the experiment, animals were sacrificed, perfused and the brains collected. Brain sections of 30-40 μm thickness were cut from frozen tissue and stained with cresyl violet, a Nissl stain. The position of the recording electrodes was determined by reference to rostral thalamus borderlines in the brain atlas (35) and visual estimation of the track of the electrodes in the brain tissue. Additionally, recording positions were determined by calculating distance above the deepest electrode position and calculating the distance below the first penetration into the tissue, compared to the known position of the electrodes below the brain surface for each recording session (expressed in μm).

---

**Supplementary Figure 1:**
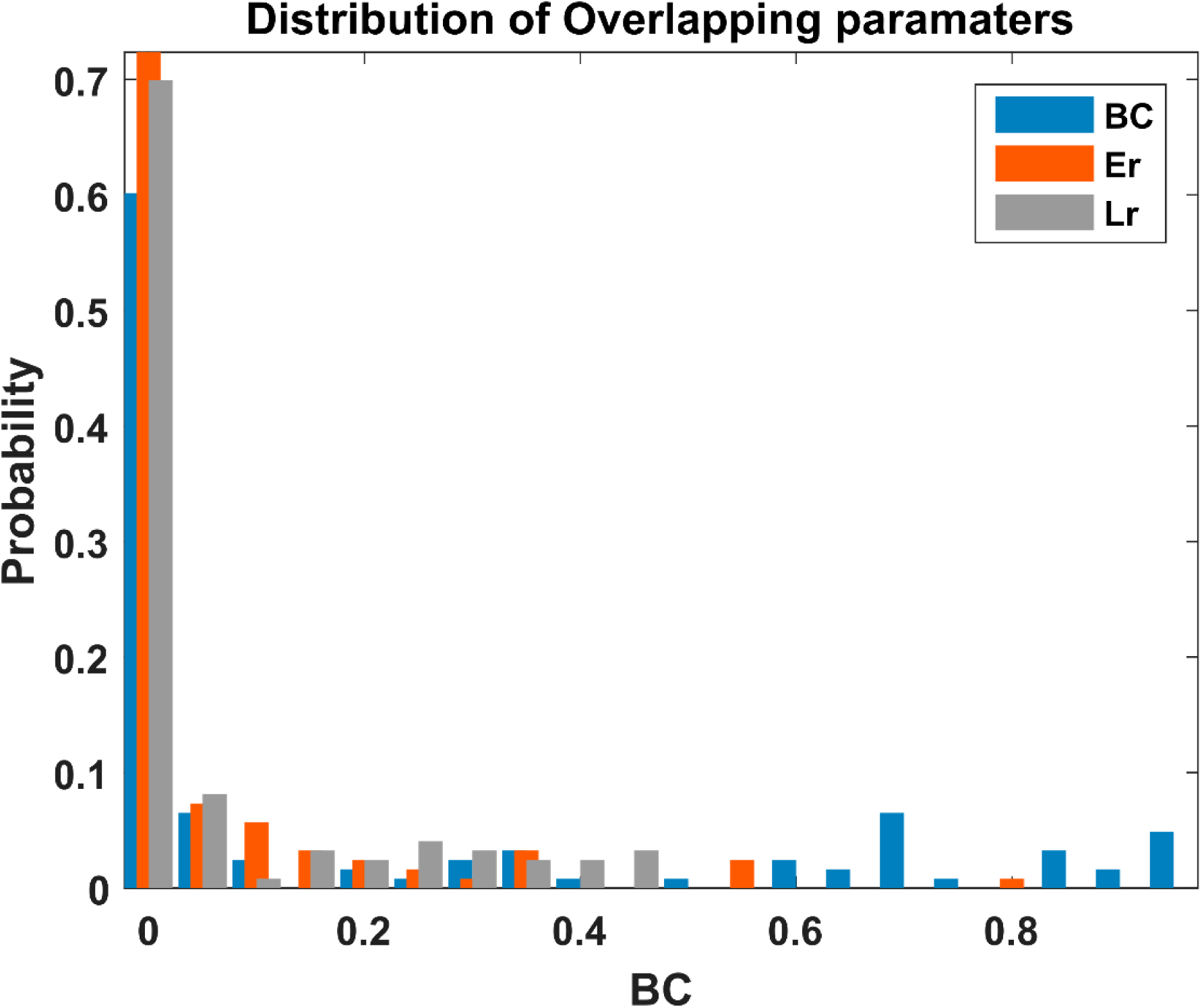
Distribution of the cluster-quality measures or measure of degree of overlap; BC= Bhattacharyya Coefficient, Er= Enclosure ratio, Lr= L-ratio.

## Acknowledgements

This work was supported by a Joint Senior Investigator Award made by The Wellcome Trust to JPA and SMOM

